# CGRP reception potentiates anxiety in an influenza A derived immune engram

**DOI:** 10.64898/2026.05.20.725748

**Authors:** Sarah K. Monroe, Benjamin A. Devlin, Ariana Vaida, Nikhita Nanduri, Hannah A. Staley, Estefany Y. Reyes, Dang M. Nguyen, Julia E. Dziabis, Aja Pragana, Seneca R. Oxendine, Mari L. Shinohara, Nicholas S. Heaton, Staci D. Bilbo

## Abstract

An immune engram is a recently described phenomenon in which neuronal populations encode functional aspects of an immune challenge. Here we investigate an immune engram arising from respiratory infection with influenza A virus, demonstrating a molecular mechanism with differential influence over behavioral and immunological aspects of the engram. We first define a cellular response to acute non-neurotropic influenza A/Puerto Rico/8/1934 (<u>PR8</u>) infection by mapping cFos+ cells and microglia morphology across brain regions. In the posterior insula, this response has an early peak at 3 days post infection. Using a cre-dependent excitatory chemogenetic system in TRAP2 mice, we capture an engram at this same region and infection timepoint. Activation of this PR8 engram results in anxiety behavior and increased transcriptional expression of cytokines in lung tissue but not spleen tissue. We further explore how pulmonary signals contribute to this PR8 engram. Using tissue-specific, cre-dependent expression of diphtheria toxin fragment in *Calca^cre^* mice, we ablate *Calca-*expressing cells including pulmonary neuroendocrine cells in respiratory tissue. Loss of *Calca*-expressing cells prevents changes in synaptic engulfment by microglia in the insula during PR8 infection without altering the cellular response to infection in pulmonary tissue. Signaling of calcitonin gene related peptide (<u>CGRP</u>), a peptide encoded by *Calca*, can be blocked with the small molecule CGRP receptor antagonist rimegepant. Using rimegepant during acute PR8 infection we again demonstrate that loss of *Calca* signaling prevents the cellular response to PR8 infection in the insula. Finally, applying rimegepant alongside the chemogenetic system in TRAP2 mice we show that CGRP receptor antagonism during engram formation prevents anxiety behavior but not peripheral gene expression changes resulting from PR8 engram activation.

## INTRODUCTION

The engram was first proposed by Richard Semon in his 1904 publication *Die Mneme*. Semon distils the idea that external experiences are internalized through physical encoding within cells—readily observed in the central nervous system (CNS) through neuronal encoding of memories. He proposed that when an organism encounters aspects of the original experience, the engram could be revived. Recently, the activity of neuronal ensembles has been shown to be sufficient to induce peripheral immunological changes in the absence of the original insulting stimuli: i.e., demonstration of an “immune engram”^1,2^. Despite excitement about this phenomenon, there is little known about the functional role or formation of immune engrams. The specificity of an immune engram to activate certain organs, the ability of an immune engram to cause complex behavioral changes, and the mechanistic underpinnings of immune engrams remain undetermined.

The lungs make up the body’s largest mucosal interface with our environment and constitute a critical interface for environmental pathogens. The connection between the lungs and brain has been well described in health and disease including viral infection^3,4^. However, the mechanisms by which respiratory infection alters the CNS are incompletely understood^5–10^. Influenza A infection has a well described effect on CNS inflammatory state and function, even in the case of non-neurotropic strains^5–9^. This literature has largely restricted its examination to inflammatory mediator impacts on the hippocampus, and hippocampal-dependent spatial memory deficits during and after influenza infection^5–7,10^. While interesting, the hippocampal response is likely insufficient to fully explain the CNS and behavioral effects of respiratory infection, especially those that may persist after the infection resolves^11^.

Here we investigate the brain’s response to respiratory viral infection with influenza A/Puerto Rico/8/1934 (<u>PR8</u>), including a resultant immune engram with functional behavioral consequences. We targeted signaling of calcitonin gene related peptide (<u>CGRP</u>), demonstrating that CGRP is a molecular signal necessary for behavioral encoding in the observed immune engram. These results expand our understanding of neuronal representation of immune stimulus, the mechanistic underpinnings, and potential means of decoupling complex aspects of an immune engram.

## RESULTS

### PR8 INFECTION CAUSES A RAPID AND TRANSIENT RESPONSE IN THE CNS

To explore which brain regions may have functional changes during influenza A infection, we began with a screen to assess cellular activity. We used the mouse adapted H1N1 influenza A virus strain A/Puerto Rico/8/1934 (<u>PR8</u>) as a model of respiratory viral infection and mapped cFos+ cells as a proxy measure for increased activity of neurons and other cells. Adult C57BL/6J mice received an intratracheal instillation of PR8 or vehicle solution. Intratracheal administration was chosen to avoid infection of the olfactory epithelium. We focused our screen at the onset of infection to ask how the CNS might initially respond to and integrate information about an emerging infection, using a non-lethal dose targeted to induce 20% weight loss at the peak of sickness. Brain tissue was collected at 2, 3, and 6 days post infection (dpi) to enable examination of tissues prior to and during onset of symptoms and systemic inflammation (Fig 1A). We applied an unbiased machine-learning based image analysis pipeline to map cFos+ expressing cells across brain regions at three coronal plates (bregma +0.25, -0.35, and -1.79). To determine which regions respond to PR8 infection, average cFos+ cell counts were compared per observed brain region between PR8 infected and control animals (Fig 1B).

**Figure 1.**
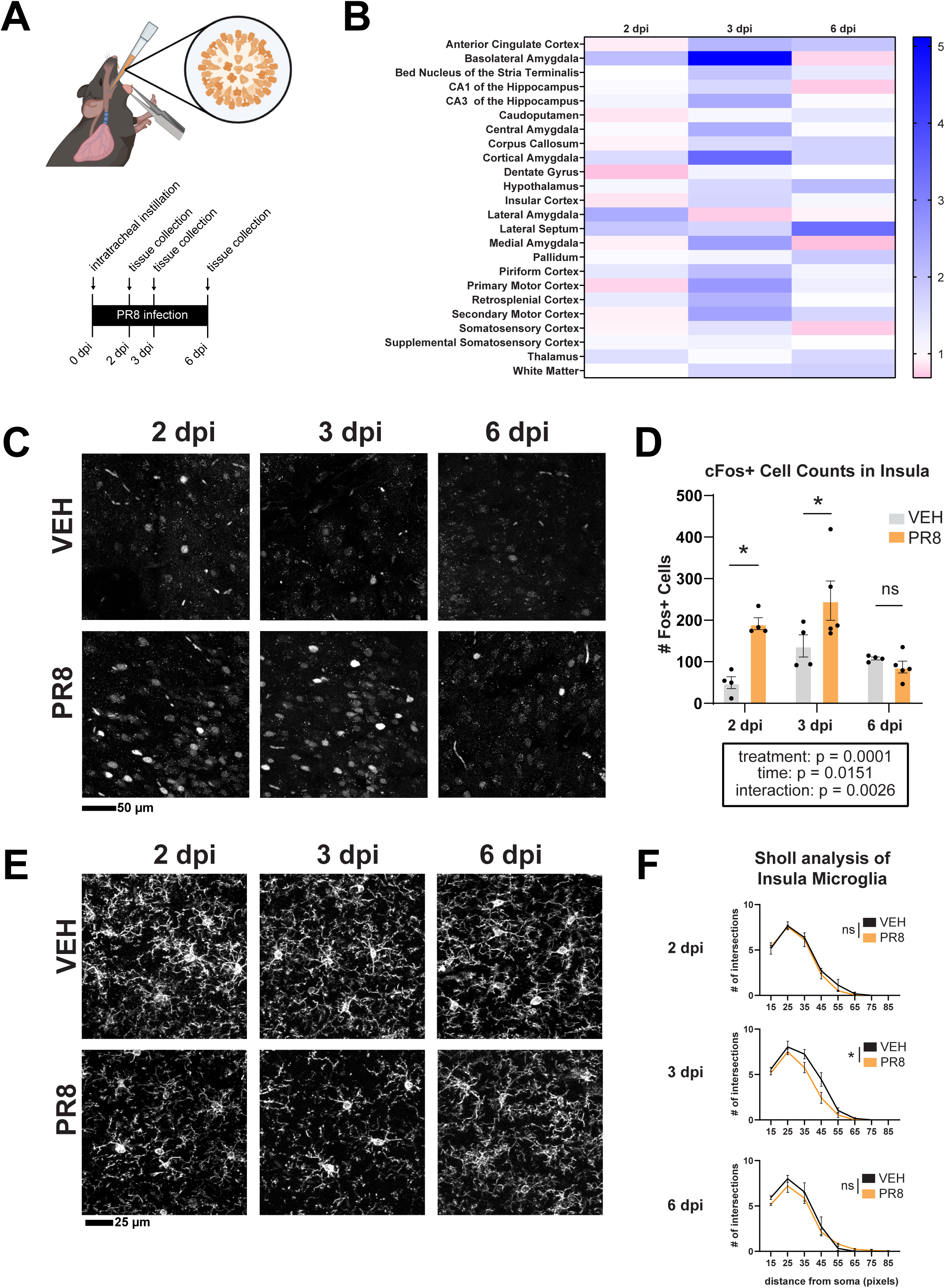
(A) Schematic of intratracheal instillation and experimental timeline. (B) Heat plot of the average fold change in cFos+ cells in PR8 infected animals compared to vehicle treated animals by brain region [n = 4]. (C) Representative images of cFos immunofluorescent staining in the posterior insula of PR8 infected or vehicle treated animals at 2, 3, and 6 dpi. (D) Quantification of cFos+ cells in the insula at 2, 3, and 6 dpi [2-way ANOVA, n = 4 – 5]. (E) Representative images of IBA1 staining in the insula of PR8 infected or vehicle treated animals at 2, 3, and 6 dpi. (F) Quantification of microglia morphology in the posterior insula at 2, 3, and 6 dpi by Sholl analysis [2-way ANOVA, n = 4 – 5].

Increased cFos+ cell counts were observed most strikingly at 3 dpi, with the highest magnitude fold change occurring in the basolateral amygdala and cortical amygdala (Fig 1B). Increases in amygdalar subregions were accompanied by a parallel response across a network of other brain regions including the nearby piriform cortex and insula, the anterior cingulate cortex, cortical motor regions, CA3 of the hippocampus, and others. Response to infection onset peaked at earlier or later timepoints in other brain regions. This dataset represents a temporally dynamic network response in the CNS to PR8 infection onset.

Given this multiregional response, we sought to focus on an area capable of integrating and coordinating infection response in the CNS. The insula—known for its role in interoception—is a connective hub which integrates information from many regions^12,13^. This includes integration of signals from amygdala and primary motor cortex, where our screen showed the greatest magnitude increase in cFos+ cells. The insula has a posterior to anterior gradient in which the posterior area is associated with integration of internal and external sensory information^12–14^, including interoceptive information about respiration and breathlessness^15–17^. Previously, the posterior insula was shown to encode an “immune engram”—an ensemble of neurons responding to and functionally encoding peripheral inflammation^2,18^. In this work, Koren *et. al.* observed a similar robust network response accompanied by functional encoding of the immune experience within the posterior insula. Given the integrative and interoceptive role of the insula, its strong cross-connectivity with the amygdala, and its demonstrated ability to encode an immune engram, we considered the posterior insula to be a candidate for integration and coordination of a multi-regional CNS response in early PR8 infection.

We examined the posterior region of the insula to determine when this area is most attuned to PR8 infection onset. In brain tissue from the same cohort, we observed significantly increased numbers of cFos+ cells during PR8 infection in the posterior insula at 2 and 3dpi, but not 6dpi (Fig 1C-D). Since PR8 infection is known to cause changes to the morphology of hippocampal microglia, we also used Sholl analysis to quantify microglial morphology across several brain regions. We observed that microglia in the posterior insula were indeed responsive to PR8 infection, which resulted in significantly reduced ramification at 3dpi (Fig 1E-F). When comparing this to other regions, microglia in the basolateral amygdala and thalamus showed no significant changes (Fig S1A-C). Microglia in the hippocampus, a region previously demonstrated to respond to PR8 infection, showed morphology changes consistent with activation, reflecting published literature^6^. Together, these data show that the posterior insula has its most striking cellular response at 3dpi.

### AN IMMUNE ENGRAM FORMED DURING PR8 INFECTION ALTERS BEHAVIOR AND PULMONARY CYTOKINE TRANSCRIPTION

We wanted to ask whether the CNS response to PR8 infection onset, which included a transient response in the posterior insula, could encode a functional immune engram. To test the functional role of neurons active during early PR8 infection, we used *Fos^tm2.1(icre/ERT2)Luo^*/J (<u>TRAP2</u>) mice and cre-dependent chemogenetic receptor expression to “capture” and reactivate ensembles of *fos* expressing neurons in healthy or infected animals. TRAP2 mice received pAAV9-hSyn-DIO-hM3D(Gq)-mCherry, encoding a cre-dependent hM3Dq excitatory DREADD, spatially targeted to the posterior insula via stereotaxic injection. At 3dpi, 4-hydroxytamoxifen (4OHT) was injected to induce recombination of the plasmid and expression of the Gq DREADD only in *fos* expressing cells (Fig 2A). Recombination of the floxed genes “captures” the active neuronal ensemble during that time window by expression of an hM3Dq receptor, enabling activity manipulation of that ensemble. Injection of the hM3Dq agonist deschloroclozapine (DCZ) activates the ensemble, enabling investigation into its functional role (FigS2A-B).

**Figure 2.**
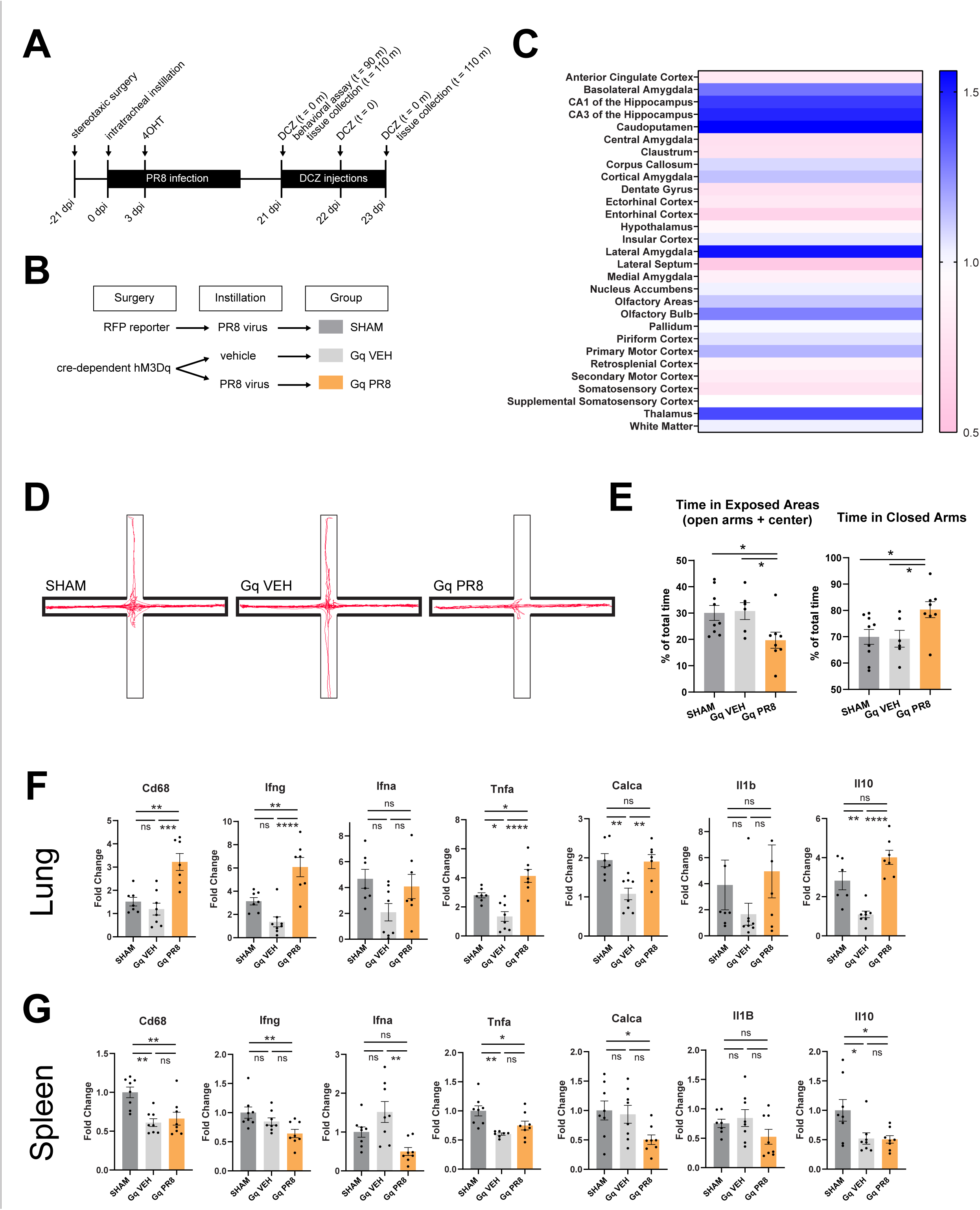
(A) Schematic of experimental timeline. (B) Schematic of experimental paradigm differentiating treatment groups. (C) Heat plot of the average fold change in cFos+ cells after DCZ stimulation in Gq PR8 animals compared to Gq VEH animals by brain region [n = 6 – 9]. (D) Representative traces tracking the center point of an experimental animal during the elevated plus maze paradigm. (E) Quantification of time spent in exposed areas (center and open arms) or closed arms as a percentage of total time in the arena [1-way ANOVA, n = 6 – 9]. (F) Fold change (ΔΔC_T_) of RNA expression for genes measured by RT-qPCR [1-way ANOVA] in lung homogenate after 3 days of DCZ stimulation [n = 6 – 9]. (G) Fold change (ΔΔC_T_) of RNA expression for genes by RT-qPCR [1-way ANOVA] in spleen homogenate after 3 days of DCZ stimulation [n = 6 – 9].

To control for the effects of the chemogenetic system and infection history, two separate control groups were used (Fig 2B). In a <u>SHAM</u> group, TRAP2 animals received an RFP fluorescent reporter plasmid instead of the cre-dependent chemogenetic system prior to PR8 infection. This group controls for any latent effects of infection that do not resolve after weeks of recovery, and any off-target effects of DCZ stimulation. In a separate <u>Gq</u> <u>VEH</u> group, the chemogenetic system was activated in healthy animals that never receive PR8. The resulting captured ensemble reflects function of *fos* expressing cells in the absence of immune stimulus, and this group controls for baseline effects of the chemogenetic system. The experimental <u>Gq PR8</u> group received the chemogenetic system, which was activated during early infection to capture the cFos+ ensemble. In Gq PR8 animals we refer to this ensemble as a “PR8 engram”. All animals recovered 21 days between intratracheal instillation and DCZ administration, enabling recovery from illness in PR8 infected groups.

We mapped cFos+ cells in Gq PR8 animals compared to Gq VEH animals to observe regions activated by DCZ stimulation (Fig 2C). We observed increased cFos+ cells after PR8 engram activation in many of the same regions that were active during acute infection including amygdalar regions. The number of cFos+ cells in PR8 infected animals increased with a greater magnitude during acute infection (up to >5 fold increase) (Fig 1B) than during PR8 engram activation (up to 1.56 fold increase) (Fig 2C). Observation of microglia morphology after DCZ stimulation was confounded by effects of the chemogenetic system, which caused a strong morphological reaction overlapping with captured neurons likely due to ongoing or latent effects of heightened neuronal activity (Fig S2C-E).

To determine whether the activated ensembles have a functional role, we assessed the behavioral response of these animals after DCZ stimulation. Given the strong response to PR8 engram activation in the amygdala, we selected an elevated plus maze (EPM) paradigm to assess anxiety-like behavior. After stimulation with one dose of the chemogenetic agonist DCZ, animals explored the EPM arena (Fig S2F). Gq PR8 animals spent significantly more time in closed arms and significantly less time in exposed areas of the maze (center and open arms) than either the Gq VEH or SHAM control groups (Fig 2D-E), demonstrating an anxiety-like behavioral phenotype. This demonstrates functional encoding of a behavioral phenotype in PR8 infected animals.

We also examined peripheral tissues of animals after ensemble stimulation by DCZ. Using RT-qPCR, we measured transcriptional expression of cytokines in lung homogenate after 3 successive days of DCZ stimulation. PR8 engram activation caused increased expression of several cytokines (*Cd68, Ifnγ, Tnfα*) in the lungs, with other genes showing increased expression attributable to infection history (*Il-10* and *Calca*) or no significant change in expression (*Il-1β*) (Fig 2G). The observed upregulation of *Cd68, Ifnγ,* and *Tnfα* could be indicative of myeloid cell activation or recruitment, known to be regulated by vagal sensory neurons within the lung^19^. Strikingly, this transcriptional pattern was not present in the spleen (Fig 2H), where PR8 engram activation caused no significant increases in measured cytokine expression, and decreased expression of *Ifnα*. While this data only captures one point in time, it portrays a distinct immunological response in pulmonary tissue rather than a homogenous systemic response to PR8 engram activation. This highlights the ability of immune engrams to encode tissue-specific information about the initial immune experience.

### MODEL ENABLES PULMONARY *CALCA*-EXPRESSING CELL ABLATION WITHOUT ALTERING PULMONARY CELLULAR REPSONSE TO PR8 INFECTION

We next investigated the pulmonary signals allowing the posterior insula to respond to PR8 infection. We targeted a sparse but critical sensory cell population in the lung epithelium: pulmonary neuroendocrine cells (<u>PNECs</u>). Like neuroendocrine cells in other tissues, PNECs are diverse signaling hubs which communicate using an array of neuropeptides^20^. Clusters of these sensory cells are directly innervated by sensory vagal neurons—notable since sensory afferents can mediate influenza-induced sickness behaviors^21^. PNECs respond acutely to immune stimuli by signaling to both pulmonary and CNS tissues^22–24^. For these reasons, PNECs stood out as likely candidates to project pulmonary immune information to the insula.

To manipulate PNECs during PR8 infection, we used a transgenic model to ablate *Calca*-expressing cells in lung tissue (Fig 3A). Adult B6.Cg-*Calca^tm1.1(cre/EGFP)Rpa^*/J (*Calca^cre^*) mice were given an AAV containing pAAV-mCherry-flex-DTA plasmid via intratracheal instillation. This method delivers a cre-dependent gene expressing diphtheria toxin fragment to epithelial cells in the trachea and lungs, while sparing cells in other critical tissues including the gut and skin. In *Calca*-expressing cells, including PNECs, the toxin fragment is expressed causing apoptosis (Fig S3A). The resulting genotypes (<u>DT-A:cre-</u>mice with intact *Calca*-expressing cells or <u>DT-A:cre+</u> mice with ablated *Calca*-expressing cells) can be compared at baseline and during infection. When comparing PR8 infection in DT-A:cre- and DT-A:cre+ mice, there were no differences in survival or weight loss through the course of infection (Fig 3B-C).This treatment significantly reduces the number of PNECs observed by histology (Fig 3D-E), and decreases *Calca* expression levels in pulmonary tissue during immune challenge with PR8 (Fig 3F).

**Figure 3.**
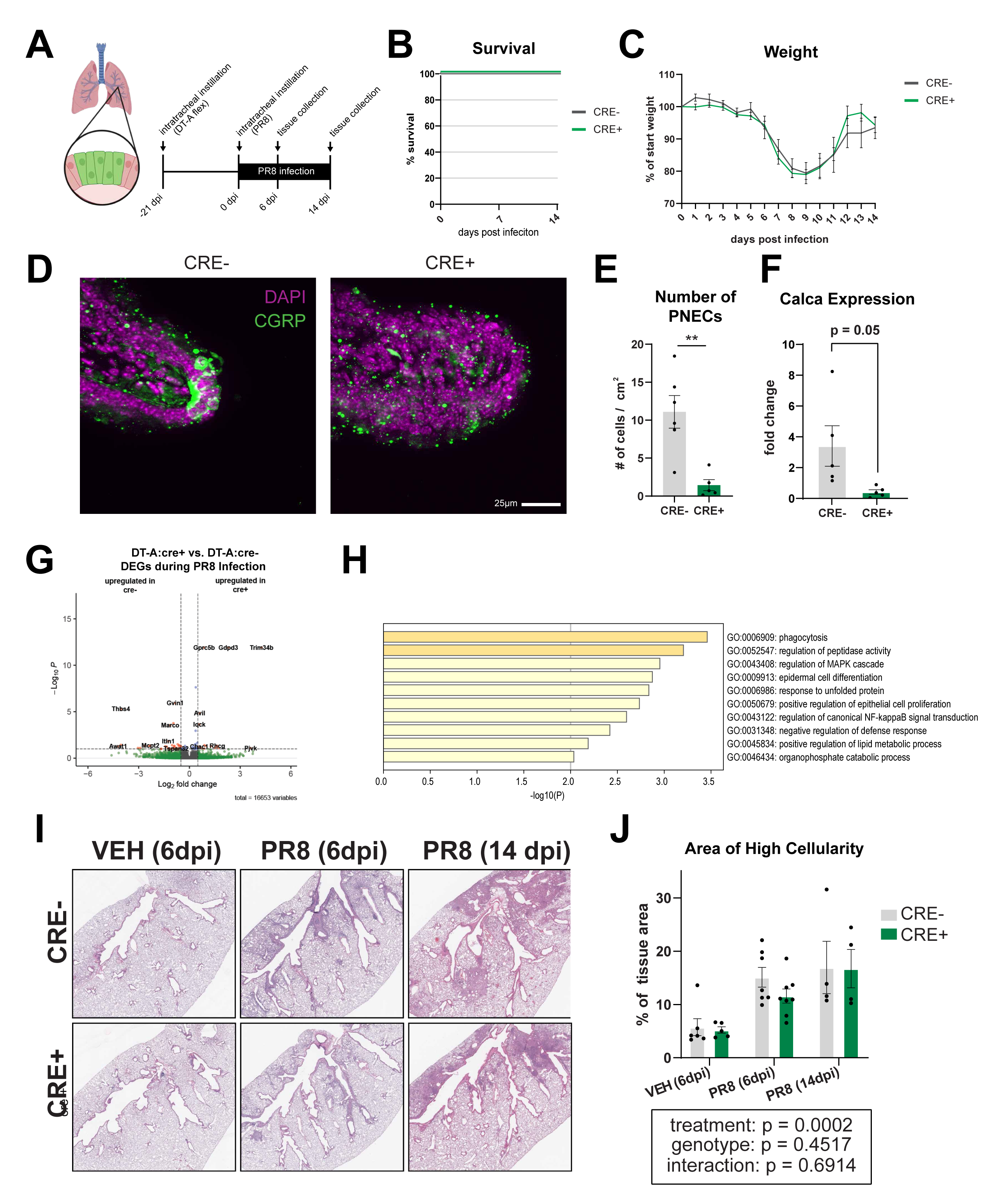
(A) Schematic of PNECs in lung tissue and experimental timeline. (B) Survival curves from 0 – 14 dpi in DT-A:cre- and DT-A:cre+ animals infected with PR8 [n = 4]. (C) Weight as a percentage of starting weight from 0 – 14 dpi in DT-A:cre- and DT-A:cre+ animals infected with PR8 [n = 4]. (D) Representative immunofluorescence images showing cell nuclei (DAPI) and PNECs (CGRP) in lung epithelial tissue of DT-A:cre- and DT-A:cre+ animals. (E) Quantification of average number of PNECs in one coronal lung section, normalized by area of lung tissue assessed [T-test, n = 5 – 6]. (F) Fold change (ΔΔC_T_) of RNA expression for *Calca* at 6 dpi in lung homogenate of DT-A:cre- and DT-A:cre+ animals infected with PR8 [T-test, n = 5]. (G) Volcano plot showing DEGs in lung homogenate of DT-A:cre- vs DT-A:cre+ animals infected with PR8 at 6 dpi; significant DEGs are shown in red [n = 5 – 6]. (H) Gene ontology of DEGs in lung homogenate of DT-A:cre- and DT-A:cre+ animals infected with PR8 at 6 dpi [n = 5 – 6]. (I) Representative images of H&E stained coronal sections of lung tissue in DT-A:cre- and DT-A:cre+ animals that are vehicle treated or infected with PR8 at 6 dpi and 14 dpi. (J) Quantification of pods as a percentage of total tissue area in coronal sections of lung [2-way ANOVA, n = 4 – 8].

To determine whether loss of *Calca*-expressing cells changed specific aspects of PR8 infection response, we began by examining the molecular landscape of the lungs. We collected tissue at 6 dpi and measured transcriptional profiles in whole lung homogenate using bulk RNA sequencing. PR8 infection caused robust upregulation of genes compared to healthy vehicle control conditions in both DT-A:cre- and DT-A:cre+ animals (Fig 3SB-E). Surprisingly however, loss of *Calca-*expressing cells minimally changed the pulmonary response to PR8 infection. When comparing genotypes, there were few significant DEGs responding to PR8 infection (Fig 3G). These results are reflected in PCA analysis, which reveals clustering of transcriptomic profiles by infection state but not by genotype (Fig S3F-G).

We also examined effects of *Calca*-expressing cell loss in DT-A:cre+ mice on PR8 infection at the cellular level. Within the lung, the cellular phenotypic infection response occurred similarly in both genotypes. We measured tissue damage response to PR8 infection using H&E histology at timepoints during and after infection. H&E histology reveals nuclei-dense pods indicating epithelial repair, which we quantified using a supervised machine learning pipeline (Fig 3I). Infection induced pods in both genotypes which continued to grow between 6dpi and 14dpi. There were no significant differences in pod area between DT-A:cre- and DT-A:cre+ animals at either timepoint (Fig 3J). We used flow cytometry to thoroughly characterize cellular response to infection in myeloid and lymphocyte populations. While infection caused an expected robust increase in immune cell populations at 6 dpi and 14 dpi, no significant differences were seen between DT-A:cre- and DT-A:cre+ animals in any measured cell population (Fig S4A-D).

Altogether, no effects were observed at the level of cellular dynamics or disease morbidity. These data indicated that while there are mild alterations in the molecular response to PR8 infection in DT-A:cre+ animals, the overall course of disease in the lung is unaltered.

### LOSS OF PULMONARY *CALCA*-EXPRESSING CELLS PREVENTS MICROGLIA RESPONSE TO PR8 INFECTION IN THE POSTERIOR INSULA

Ultimately, we sought to determine whether loss of *Calca-*expressing cells in this model changed the CNS response to PR8 infection. Previously we observed a cellular response to PR8 infection in the posterior insula. We chose to focus on this same brain region, this time examining microglia at a functional level.

Engulfment of cellular materials or compartments is a more robust indicator of microglia function compared to morphology alone. Microglia often alter their engulfment of synapses in response to changes in neural activity (Shafer *et. al.*, 2012; Devlin *et. al.*, 2025). To assess whether microglia altered their interactions with synapses in this model, we created digital 3D reconstructions of microglia from high resolution confocal images taken in the posterior insula from the same animals examined above (Fig 4A). We immunohistochemically stained to observe microglia morphology, their lysosomes, and either vGlut2 or vGat to label excitatory or inhibitory pre-synaptic markers, respectively. Using IMARIS reconstructions, we quantified phagocytosed synaptic material within the lysosomes of these cells as a percentage of total cell volume. Infection caused a reduction in both vGlut2 and vGat engulfment during PR8 infection in DT-A:cre- mice; this effect was prevented in DT-A:cre+ mice (Fig 4B - E). These data again implicate the posterior insula as a brain region that is responsive to PR8 infection, including functional changes in microglia. It also suggests that pulmonary signals, specifically from *Calca*-expressing cells, are enabling a cellular response to PR8 infection in the posterior insula.

**Figure 4.**
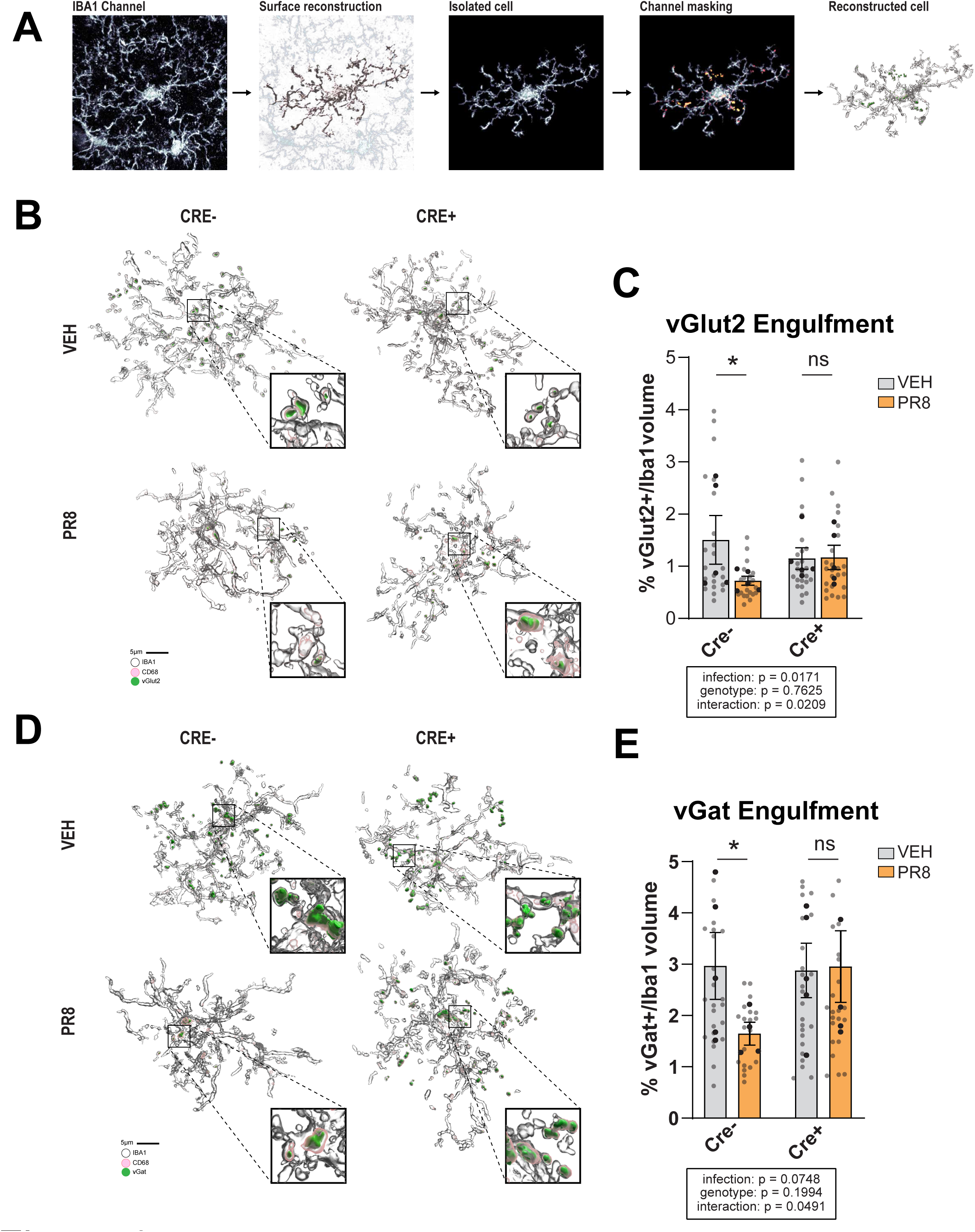
(A) Schematic representation of the 3D reconstruction process, showing proregression from the raw image to a reconstruction of a cell surface, isolation of one cell surface, masking of the lysosomal channel within that cell and engulfed material within those lysosomes, and final reconstruction output. (B) Representative 3D reconstructions made from immunofluorescence images of microglia (IBA1+), their lysosomes (CD68+) and engulfed vGlut2+ material within lysosomes after PR8 infection in DT-A:cre- and DT-A:cre+ animals. (C) Quantification of engulfed vGlut2+ material in the lysosomes of microglia reconstructions expressed as a percentage of total cell volume [2-way ANOVA, n = 22 – 25]. (D) Representative 3D reconstructions made from immunofluorescence images of microglia (IBA1+), their lysosomes (CD68+) and engulfed vGat+ material within lysosomes after PR8 infection in DT-A:cre- and DT-A:cre+ animals. (E) Quantification of engulfed vGat+ material in the lysosomes of microglia reconstructions expressed as a percentage of total cell volume [2-way ANOVA, 20 – 25].

### RIMEGEPANT PREVENTS CELLULAR RESPONSE TO PR8 INFECTION IN THE POSTERIOR INSULA

Given that the microglia response during acute PR8 infection is sensitive to signals from pulmonary *Calca*-expressing cells, and that this influence occurs in the same region where we captured a functional PR8 engram, we sought to determine whether signals from *Calca*-expressing cells could be influencing the PR8 engram itself. Since it is not possible to combine the functionality of cre-based TRAP2 mice and *Calca^cre^* mice, we opted to pharmacologically inhibit a key signaling pathway engaged by PNECs.

PNECs produce many important neuropeptides in the lung, and the loss of *Calca*-expressing cells in our DT-A:cre+ mice may be influencing many aspects of PNEC signaling including CGRP, GABA, serotonin, vagal signals, and more. CGRP, a peptide encoded by *Calca*, is known to induce anxiety^25^, a behavior encoded within the PR8 engram we observed. We used a targeted pharmacological approach to block CGRP signaling without disturbing other molecular or vagal signaling from PNECs by administering rimegepant, a small molecule CGRP receptor antagonist. Recently, rimegepant has also been demonstrated to alter asthmatic response in a mouse model when peripherally administered^26^, mirroring experimental outcomes after loss of *Calca-*expressing cells in murine models of asthma^27^. Given parallel outcomes in this literature, we applied rimegepant in TRAP2 mice as a more targeted proxy for *Calca*-expressing cell ablation.

Before exploring effects of rimegepant on the PR8 engram, we first asked how rimegepant affects the cellular response to PR8 in the posterior insula during acute infection. We infected adult C57BL/6J animals with PR8 or administered a vehicle control, then administered rimegepant or vehicle solution every 24 hours at 1 and 2 dpi. We collected tissues 6 hours after the final dose of rimegepant. In vehicle treated animals we observed an increase in cFos+ cells (Fig 5A-B) and more amoeboid microglia morphology in the posterior insula after PR8 infection (Fig 5C-D), consistent with data discussed above (Fig 1D-G). In rimegepant treated animals, there was no longer any increase in cFos+ cells in the posterior insula after PR8 infection (Fig 5A-B). Additionally, the magnitude of morphological changes in microglia after PR8 infection in this region were reduced (Fig 5C-D). This effect aligns nicely with the loss of changes in synaptic engulfment seen in the *Calca*-expressing cell ablation model above (Fig 4B-E). In both models, loss of CGRP signaling reduces the cellular response to PR8 infection in the posterior insula.

**Figure 5.**
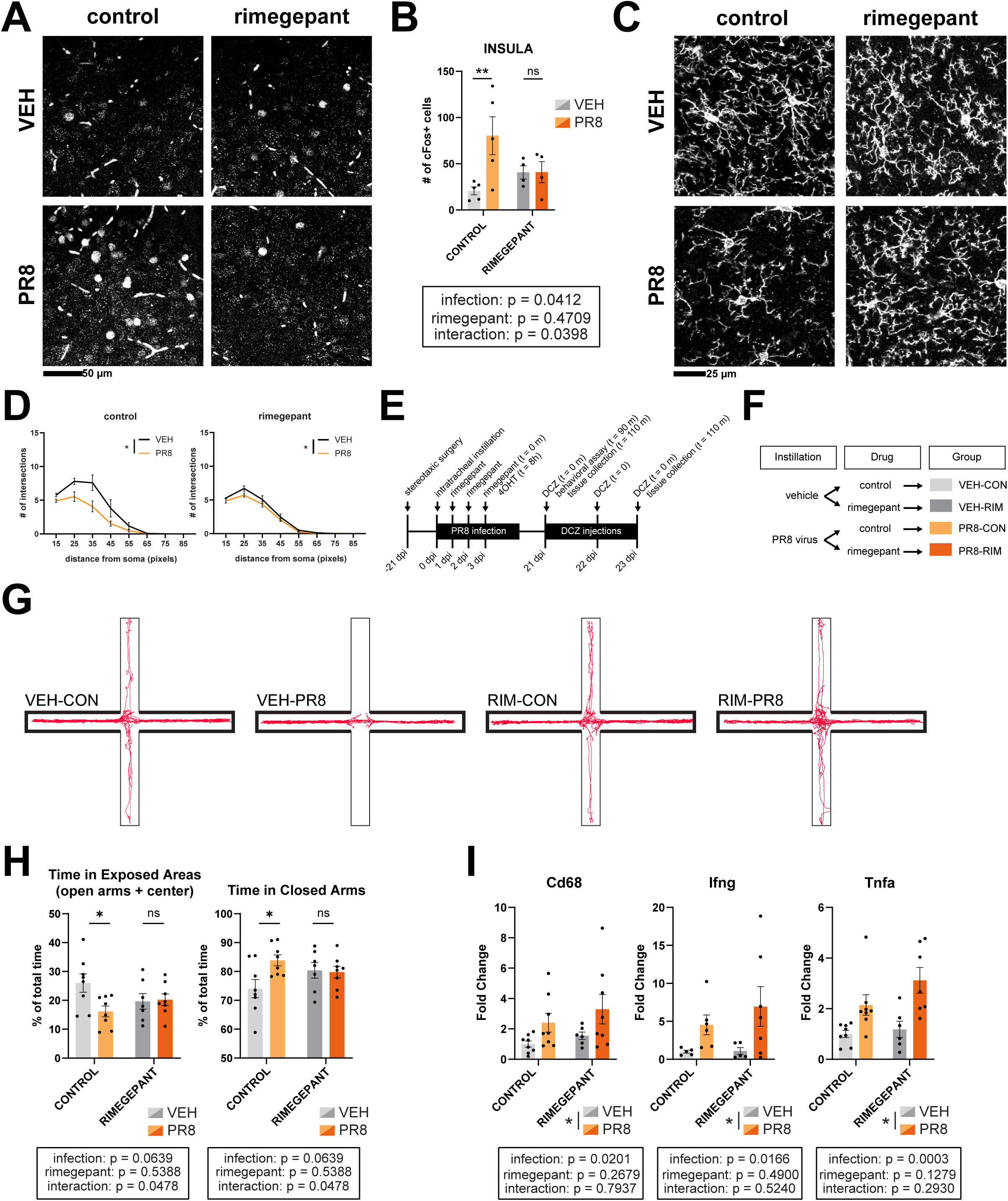
(A) Representative images of cFos immunofluorescent staining in the posterior insula. (B) Quantification of cFos+ cells in the posterior insula [2-way ANOVA, n = 4 – 5]. (C) Representative images of IBA1 immunofluorescent staining in the posterior insula. (D) Quantification of microglia morphology in the insula by Sholl analysis [2-way ANOVA, n = 4 – 5]. (E) Schematic of experimental timeline. (F) Schematic of experimental paradigm differentiating treatment groups. (G) Representative traces tracking the center point of one experimental animal during the elevated plus maze paradigm. (H) Quantification of time spent in exposed areas (center and open arms) or closed arms as a percentage of total time in the arena [2-way ANOVA, n = 7 – 8]. (I) Fold change (ΔΔC_T_) of RNA expression for genes measured by RT-qPCR [2-way ANOVA, n = 5 – 8] in lung homogenate after 3 days of DCZ stimulation.

### RIMEGEPANT ELIMINATES BEHAVIORAL ASPECTS OF A RESPIRATORY INFECTION ENGRAM

To finally test whether CGRP reception influences the observed PR8 engram, we administered rimegepant as an intervention in our model of engram capture in TRAP2 mice (Fig 5E). Adult TRAP2 mice were stereotaxically injected in the posterior insula with the cre-dependent hM3Dq encoding plasmid as before. After recovery, animals received a PR8 infection or vehicle instillation and were administered either rimegepant or a vehicle solution every 24 hours until ensemble capture by 4OHT injection at 3dpi. Infection then proceeded normally, with engram reactivation occurring at the same post-recovery timepoint as in prior experiments. This resulted in four treatment groups varying by rimegepant and infection status (Fig 5F). The VEH-CONTROL and PR8-CONTROL groups are analogous to the Gq-VEH and Gq-PR8 groups above, capturing and activating a neuronal ensemble in healthy animals or a PR8 engram. The VEH-RIMEGEPANT group controls for baseline effects of rimegepant, and the PR8-RIMEGEPANT group examines how blocking CGRP reception influences the PR8 engram. We repeated EPM and peripheral cytokine gene expression measurements in these animals after DCZ stimulation.

Elevated plus maze in VEH-CONTROL and PR8-CONTROL animals again demonstrated that PR8 engram activation significantly increases anxiety-like behavior compared to healthy ensemble activation (Fig 5G-H). Strikingly, in groups receiving rimegepant the behavioral outcome was the same regardless of infection status during engram capture. This indicated that behavioral aspects of the PR8 engram are lost when CGRP reception is blocked, matching loss of microglia response in the posterior insula after *Calca*-expressing cell ablation. These data point to CGRP as a molecular signal which enables engram encoding of anxiety during PR8 infection.

In the lung tissue of the same animals, PR8 engram activation again caused increased expression of genes related to myeloid cell activation (*Cd68, Ifnγ, Tnfα*) (Fig 5I) in concordance with prior experiments (Fig 2G). Unlike anxiety behavior, this gene expression pattern was retained in rimegepant-treated animals. These data indicate that CGRP reception is not necessary to encode this pulmonary aspect of PR8 engram activation.

## DISCUSSION

In capturing and reactivating neuronal ensembles activated during acute infection, we explored the existence and functional consequence of a pulmonary immune engram. We found that both anxiety and pulmonary gene expression can be altered by engram activation. To our awareness, while other states including lethargy and appetitive suppression have been observed after immune engram activation^1^, this is the first demonstration of an emotional behavior arising from an immune engram. The differences in gene expression changes between lung and spleen tissue during PR8 engram activation further show that immune engrams can encode tissue-specific information across the periphery. Lastly, we showed that CGRP signaling influences behavioral but not immunological aspects of this pulmonary immune engram. This work advances our understanding of immune engram formation and function.

Mapping of cFos+ cells revealed a temporally dynamic network response during PR8 infection onset. It is important to note that while certain regions displayed a greater magnitude of increase in cFos+ cell counts, this is just one measure of CNS function which does not comprehensively reflect biological relevance.

During infection, the cellular response to PR8 infection in the posterior insula is sensitive to CGRP signaling. Microglia in the posterior insula showed a transient response to PR8 infection, developing more amoeboid morphology at 3 dpi when cFos+ cell counts peak. This morphological response could be induced by purinergic reception of increased neuronal activity in the surrounding tissue, or through sensation of a circulating peripheral signal that crosses the blood-brain barrier. An additional possibility is that a transiently “leaky” blood-brain barrier enables microglia in the posterior insula to respond to a peripheral signal at 3dpi that would not typically reach these cells^28^. This situation opens the possibility of posterior insula microglia responding directly to peripheral CGRP through CALCRL or RAMP1 receptors^29^. Regardless of whether the signal is direct or indirect, loss of CGRP signaling affects the cellular response to PR8 infection in the posterior insula. Both *Calca*-expressing cell ablation and rimegepant reduced the ability of microglia in the posterior insula to respond to PR8 infection.

Our model of ensemble capture and activation represents a functional immune engram in animals that experienced a PR8 infection. While the ensemble we manipulated was centered on the posterior insula, it innervates and activates the amygdala, motor cortex, piriform cortex, and other areas where PR8 infection itself causes a cellular response. Engram capture and manipulation did not influence the posterior insula in isolation but instead engaged a larger network response across the brain.

This engram encoded a behavioral phenotype and altered transcriptional function, demonstrating the broad reaching potential of immune engrams. The gene expression changes we observed after PR8 engram activation were distinct between the lungs and spleen, indicating that the PR8 engram may encode tissue-specific immune responses. Together, these data demonstrate a PR8 engram which encodes complex information relating to the original immune event.

Loss of CGRP reception by administration of rimegepant was able to influence the PR8 engram, preventing encoding of anxiety behavior. In the same animals, rimegepant failed to alter upregulated expression of cytokines in lung tissue resulting from PR8 engram activation. These data suggest that behavioral and immunological aspects of immune engrams can be distinctly encoded and decoupled. The observed PR8 engram involves activation and suppression of activity across a broad network of brain regions, so it is unsurprising that this CNS response involves multiple mechanistic pathways. Whether microglia pruning or functional activity is necessary for anxiety encoding in the PR8 engram remains to be seen and is the planned future direction of this work.

To speculate on the function of the observed PR8 engram, if an organism could recognize cues of an immune threat in its environment (e.g. an infected peer) and activate the observed engram, then the resulting stimulation of cytokine transcription could mount a pre-emptive defense against the immune threat. Anxiety behavior could be beneficial in threat avoidance and further aid in survival. Within human society there are situations where approaching environmental cues of sickness are necessary or desirable. For example, healthcare providers engage with cues of sickness as part of their daily work, particularly in the context of contagious disease outbreak. In these scenarios, decoupling anxiety from protective immune function would be highly desirable. Future work can uncover more about the functional role of engrams, including which environmental cues activate an immune engram and whether behavioral or immunological effects of engram activation can reduce probability of future infection.

## METHODS

### Animals

Animal care and procedures were conducted in accordance with the NIH Guide to the Care and Use of Laboratory Animals and approved by the Duke University Institutional Animal Care and Use Committee (IACUC). Animals were group housed in a standard 12:12-hour light–dark cycle. Animals had continuous access to chow, water, corn cob bedding, and at least 2 forms of enrichment. The C57BL/6J (wild type) mouse line was obtained from The Jackson Laboratory, stock no. 000664. The B6.Cg-*Calca^tm1.1(cre/EGFP)Rpa^*/J (*Calca^cre^*) mouse line The Jackson Laboratory, stock no. 033168. The *Fos^tm2.1(icre/ERT2)Luo^*/J (TRAP2) mouse line The Jackson Laboratory, stock no. 030323. Once obtained, transgenic mouse lines were bred in house with C57BL/6J females to produce experimental offspring. Genotyping of transgenic animals was conducted on tail or ear tissue DNA using polymerase chain reaction in accordance with published protocol from The Jackson Laboratory for each mouse strain. Experiments were performed on both male and female animals, with no statistically significant sex differences observed when considering sex as a variable. Adult animals used in these experiments are defined as age P60 – P115, with ages distributed evenly across treatment groups within each experiment.

### Intratracheal viral instillations

Animals were anesthetized using 4.5% inhalational isoflurane in O2 to achieve surgical plane 3, at which point isofluorane was lowered to 2%. Animals were then suspended supine on an 80° incline. Blunt tools were used to extend the tongue and light pressure was applied to block the nares. A pipette was used to administer 50 µL of liquid to the trachea, then animals were held in position for 60 breaths. Animals were monitored until full recovery.

### Viral handling

Influenza A: A/Puerto Rico/8/1934 (PR8) viral stock was purified from inoculated chicken embryos and titer was measured by plaque assay. Stocks were thawed over ice and diluted with sterile PBS to create working aliquots of 1000 PFU/µL. Aliquot titer was reconfirmed with plaque assay. Aliquots were thawed on ice immediately prior to instillation and diluted to 3000 PFU per 50 µL dose in a 0.9% sterile saline vehicle. Instillation was performed in a BSL2 hood. Animals underwent daily health monitoring including weight measurement and observational assessment inside a BSL2 hood.

pAAV6-mCherry-flex-DT-A: pAAV-mCherry-flex-DTA plasmid was obtained from AddGene (Plasmid #58536), packaged at the Duke University Viral Vector core, aliquoted and stored at -80°C. Viral aliquots were thawed on ice immediately prior to instillation and diluted to ##vg per 50 µL dose in 0.9% sterile saline vehicle.

pAAV9-hSyn-DIO-hM3D(Gq)-mCherry: Packaged plasmid was obtained from AddGeen (plasmid # 44361) and stored at -80°C. Viral aliquots were thawed on ice and kept over ice for the duration of stereotaxic surgery.

pAAV5-hSyn-mCherry: Packaged plasmid was obtained from AddGene (plasmid #114472) and stored at -80°C. Viral aliquots were thawed on ice and kept over ice for the duration of stereotaxic surgery.

### Chemogenetic manipulation

Animals underwent bilateral stereotaxic surgery to inject pAAV9-hSyn-DIO-hM3D(Gq)-mCherry or pAAV5-hSyn-mCherry into bregma coordinates AP, −0.35 mm; ML, 4.0 mm; DV, −3.83 mm. Animals were anesthetized using 4% inhalational isoflurane in O2 to achieve surgical plane 3, at which point isoflurane was lowered to 2.5%. Animals received a subcutaneous injection of ketoprophen analgesic (5mg/kg), a subcutaneous injection of 0.25% bupivacaine HCL under the incision site, and periodic subcutaneous injections of 0.9% sterile saline throughout surgery. Animals were fitted to a Stoelting stereotaxic apparatus for the duration of the surgery. Skin was shaved prepped with betadine prior to surgical incision on the scalp. Bregma was identified and bilateral entry holes were drilled a measured distance away above the injection area. 1×10^13^ vg were injected at 0.7 µL/min using a Hamilton 7001Kh 1µL syringe in each hemisphere. Needles were kept in place for 5 minutes after the cessation of injection. Animal incision was dressed appropriately, and animals were monitored until full recovery. Animals were monitored and administered ketoprophen analgesic (5mg/kg) each 24 hours following surgery. At 21 days after surgery, animals underwent Influenza A: A/Puerto Rico/8/1934 instillation and monitoring as described above. At 3 days post infection, animals received an I.P. injection of 75 mg/kg 4-hydroxytamoxifen (Millipore Sigma; H6278-50MG) mixed overnight at 37 degrees C in 16.7% EtOH in corn oil. At 21 days post infection, or at days 21 – 23 post infection, animals received an I.P. injection of 0.1 mg/kg deschloroclozapine (DCZ) (Bio Techne, Cat. No. 7193) dissolved in 1% DMSO in 0.9% sterile saline. Animals underwent behavioral assays at 90 minutes after initial DCZ injection. Animals were sacrificed 110 minutes after DCZ injection at either 21 or 23 days post infection.

### Elevated Plus Maze

Animals were placed in the behavioral testing room for at least 60 minutes prior to testing. On a given day, males were tested prior to females with no opposite-sex animals present in the room during any testing or acclimation period. One at a time, animals were set in the center of an elevated plus maze consisting of a symmetrical plus-shaped platform with four 5 cm wide x 36 cm long arms, two of which having 15 cm tall walls, and a 5 cm x 5 cm center area. Animals were allowed 10 minutes to explore the maze, and a ceiling mounted webcam was used to create video recordings of the tests. The maze was cleaned between each animal with 70% ethanol and allowed to dry prior to placement of the next animal. Location in video files was tracked and analyzed using Ethovision (Nodulus) to identify which portion of the maze an animal was in at each frame of the recording. “Percent Time in Closed arms” was calculated by (time in closed arms / time of total recording duration) *100. “Percent Time in Exposed Areas” was calculated by ([time in open arms + time in center] / time of total recording duration) *100.

### Rimegepant

Rimegepant (BMS-927711; CAS No. 1289023-67-1) was obtained from MedChemExpress (Cat. No.: HY-15498) and stored at 4°C. Rimegepant was weighed and prepared for *in vivo* use according to manufacturers instructions by adding the following solvents sequentially: 5% DMSO, 40% PEG300, 5% Tween-80, 50% Saline. Control solution was prepared in tandem using the same solvents without any Rimegepant. Animals were administered 10 mg/kg Rimegepant or vehicle control solution by I.P. injection between ZT 2 and ZT 4. In experiments where tissue collection occurred on the same day, animals rested in their home cage for 6 hours between injection and collection.

### Tissue collection

After CO2 euthanasia, animals were transcardially perfused with ice-cold saline. When used for molecular work, lung and spleen tissues were removed and flash frozen on dry ice and stored at -80 °C. When used for histology, lungs were inflated via intratracheal needle with a 1:1 mixture of 8% PFA and OTC, tied below the needle entry point with suture thread, clipped from connective tissue, and drop fixed for 48 hours in 4% PFA. Brains were removed and drop fixed for 48 hours in 4% PFA. After PFA fixation, tissues used for immunohistology were placed in a 30% sucrose in 1X PBS solution with 0.1% sodium azide for at least 72 hours at 4°C. Alternatively, tissues used for hematoxylin and eosin histology were moved to 70% ethanol and stored briefly at 4°C.

### Hematoxylin and eosin histology and quantification

Lung tissues were transferred to Histowiz (Long Island City, NY) for paraffin embedding, sectioning, staining, and imaging. Images were quantified using QuPath cell identification and pixel classifiers to label nuclei, segment tissue from background, segment cellularly dense tissue from typical alveolar tissue, and measure segmentation areas.

### Immunohistochemistry

Tissues were flash-frozen in 2-methylbutane pre-chilled on dry-ice, and stored at -80°C until cryosectioning at 40µm into tubes of cryoprotectant. Sectioned tissue was stored at -20°C until use. Free floating slices were pulled from sectioned material and washed 3 times in PBS (washes were 10 minutes each, with tissue at room temperature and shaking at 50 rpm). Tissue was then shaken at 50 rpm at room temperature in blocking solution (10% normal serum, 0.3% Triton-X 100 and 1% H2O2 in PBS) for at least 1 hour. Tissue was then moved to tubes of primary antibody solution (2% normal serum, and 0.3% Triton-X 100, and primary antibodies [see Table 1] in PBS) at stored at 4°C overnight. Tissue was then washed 3 times in PBS as before and moved to tubes of secondary antibody solution (2% normal serum, and 0.3% Triton-X 100, and secondary antibodies [see Table 1] in PBS) to be shaken at 50 rpm for 2 hours at room temperature. Tissue was washed a final 3 times in PBS, wet mounted in 5% BSA onto gelatin-subbed (0.1% gelatin and chromium potassium sulfate) glass slides, set with vectashield PLUS mounting medium containing DAPI (Vector Laboratories; H-2000-2) under a glass coverslip, and sealed with acrylic nail polish.

**Table 1.** Immunohistochemistry antibodies.

| Antigen | Brand | Catalog Number | Concentration |
| --- | --- | --- | --- |
| CD68 | BioLegend | 137002 | 1:1000 |
| cFos | Millipore | ABE457 | 1:1000 |
| CGRP | Millipore | C8198 | 1:200 |
| IBA1 | Novus | NB100-1028 | 1:1000 |
| IBA1 | Synaptic Systems | 234 009 | 1:1000 |
| RFP | Synaptic Systems | 390 004 | 1:1000 |
| vGat | Synaptic Systems | 131 004 | 1:1000 |
| vGlut2 | Synaptic Systems | 135 404 | 1:1000 |

### Imaging

Images were acquired on an Olympus FV3000 inverted confocal. For brain slice mapping, images were taken using a 20x objective (in figure 1 experiments) or a 10x objective (in all other experiments) to tile scan entire slices with each tile in a single Z plane and focus adjusted across the area of the slice. For 2-D Sholl analysis and cFos+ cell counting, images were taken using a 20x objective to tile scan brain regions in multiple z planes. For PNEC quantification, images were taken using a 4x objective to tile scan entire tissue sections, For 3-D microglia reconstruction, images were taken using a 63x oil mount objective to image several unique fields of view centering on at least one microglia in multiple Z planes. Laser channels and intensity were set per experiment based on signal intensity and kept consistent across all images within a given experiment.

### Registration and mapping of cFos+ cells and viral expression

Tilescans were stitched using Olympus FluoView software and the QUINT workflow^30^ was used for atlas registration, incorporating Ilastik^31^ for automated Fos+ cell counting. First, the two channels were separated (cFos or RFP and DAPI), automatically thresholded, and down sampled in FIJI using custom macros. Resolution was reduced to ≤3000 px in all dimensions to accommodate efficient segmentation performance and ensure pipeline compatibility. For cFos mapping, the cFos channel was processed in Ilastik using the “Pixel and Object Classification” workflow to generate object-level segmentations. The DAPI channel was first processed using DeepSlice^32^ to get preliminary registration to the mouse Allen Brain Atlas. From there, registrations were manually checked and adjusted for correctness in QuickNII^33^. Final section registrations were saved and combined with Ilastik Fos object outputs using the NUTIL quantifier^34^ loaded with custom regions annotated from the Allen Brian Atlas 2017. Per section NUTIL outputs were post-processed using custom python scripts. First, output data including “object count” and “region pixels” were combined with metadata for each section. Regions that were lowly represented in a section (<20,000 total pixels) were filtered out and object counts were normalized to remaining region area by simple division (i.e. obj count / region area) to acquire an object density measurement. From there, a scaling factor was calculated to convert pixels to microns using image metadata acquired after down sampling resolution (e.g. 0.3175 pixels per micron). This scaling factor was squared, and the region-normalized object counts were once again divided by their scaling factor to obtain an object/micron^2^ count. These values were multiplied by 10,000 to get object counts per 10,000micron^2^ area of a given brain region. Data was then normalized such that the control mean was 1, and regions were sorted based on the difference between the experimental group mean and control mean.

### Quantification of cFos+ cells in individual brain regions

Maximum intensity projections were made from Z-stack images, and the cFos staining channel was separated. The machine-learning software ILASTIK was trained per experiment on a subset of images to automatically segment and count cFos+ cells across the entire image set. cFos+ cell counts were normalized to brain region area in images where brain regions took up variable portions of the field of view.

### 2-D sholl analysis

Maximum intensity projections were made from Z-stack images, and the appropriate IBA1 staining channel was separated. Images were scrambled and treatment group was blinded before experimenters manually selected representative microglia cells within an image using FIJI. 6 – 10 cells were used per animal, using consistent cell numbers within a given experiment. Cells were selected bilaterally across hemispheres of a given animal. The machine-learning software ILASTIK was trained per experiment on a subset of images to automatically segment pixels of microglia cells from background pixels. A custom python script based on the FIJI sholl analysis plugin was used to select the center of each cell, skeletonize the processes, overlay concentric rings, and calculate number of intersections between each ring and the cell processes. Number of intersections at a given ring distance was averaged between all cells to create a single value per animal. Values per animal were averaged within a treatment group. Custom analysis scripts are available on GitHub at: https://github.com/bendevlin18/sholl-analysis-python

### PNEC quantification

Coronal sections of 40µm lung tissue were direct mounted and stained for CGRP protein using immunofluorescence. Tissue was blinded and entire slices were systematically searched to hand count CGRP+ cells. Tile scans of entire tissue sections were taken at 4x in a single Z plane. A QuPath pixel classifier was trained to identify lung tissue then used to calculate area of lung tissue. CGRP+ cell counts were normalized to tissue area to calculate absolute number of cells per cm^2^ for direct comparison between animals. Representative images were taken using a 20x objective.

### 3-D microglia reconstructions

Z-stack images were converted to IMARIS files (v9.5), and the surfaces tool was used to render the IBA1 channel into a 3-D volume. Area masking of the IBA1 surface was used to restrict the CD68 signal; this restricted CD68 signal was rendered with the sufaces tool. Area masking of the CD68 surface was used to restrict synaptic material signal (vGlut2 or vGat), and this restricted signal was rendered with the surfaces tool to make a final area of engulfed synaptic material. IBA1 surfaces for each single cell were manually selected, and the CD68 and synaptic surface volumes within that cell were calculated. Percentage of microglia engulfing synaptic material was calculated as (total volume of [vGlut2 or vGat] restricted to CD68 within IBA1 volume of a cell / total IBA1 volume of the same cell)*100. Within each animal, at least 6 cells were measured. Values per animal were averaged within a treatment group.

### RNA extraction

Lung or spleen tissues were homogenized in trizol using a Bead Bug 6 Homogenizer (Benchmark Scientific) by shaking with a 3 mm zirconium bead 3 times at 2500 rpm for 30 seconds. Chloroform was added to the homogenate (1:5 with trizol), then the solution was vortexed for 2 minutes at 2000 rpm, allowed to rest at room temperature for 3 minutes, and spun at 12,900 RCF for 15 minutes at 4°C in a centrifuge. The aqueous phase was separated, mixed 1:1 with isoproponal, vortexed for 1 minute at 2000 rpm, allowed to rest at room temperature for 10 minutes, and spun at 12,900 RCF for 10 minutes at 4°C in a centrifuge to precipitate an RNA pellet. Pellets were washed twice with 75% ice-cold ethanol followed by 5 minute centrifugation at 7,500 RCF and 4°C, then dried and resuspended in RNAse-free water.

### Bulk RNA sequencing and analysis

Lung RNA sample concentration was determined by nanodrop spectrophotometry. Samples were transferred on to MedGenome (Foster City, California) for library preparation and sequencing. Raw fastQ files were aligned to Mus Musculus GRCm38 mm9 using STAR (v2.7.5c) and featurecounts (v1.6.3) on the Duke Compute Cluster with custom bash scripts. Genes were filtered and only included in analysis if they were present at 10 counts in at least 4 samples. Differentially expressed genes were calculated using DESeq269 in R 4.4.0. All analysis scripts are available on GitHub at: https://github.com/bendevlin18/IL34blocking_seq_analysis_2024.git. Data is available in the GEO database under accession number GSE330855.

### RT-qPCR

Lung RNA sample concentration was determined by nanodrop spectrophotometry. 1 mg of RNA was used per sample for cDNA synthesis in accordance with QuantiTect Reverse Transcription Kit protocol (Qiagen; Cat. No. 205311). Three negative control samples were created during cDNA synthesis: one missing reverse transcriptase, one missing RNA template, and one that is only water. For the qPCR reaction, a master mix was made (6.5 µL SYBR (Qiagen; Cat. No. 330509), 1 µL each of forward and reverse primers (see Table 2), and 3.5 µL nuclease free water per well) and distributed in a 96 well plate alongside 1 µL of cDNA reaction product per well. Plates were sealed and vortexed at 2,000 rpm for 1 minute followed by brief centrifugation using a table top centrifuge. Plates were immediately run on a QuantStudio 3 Real-Time PCR System (Fisher Scientific). ΔΔCt values were calculated in reference to 18S Ct values from the same cDNA samples. Fold change was calculated by normalizing ΔΔCt values to the average ΔΔCt value of a control group in each given experiment.

**Table 2.**
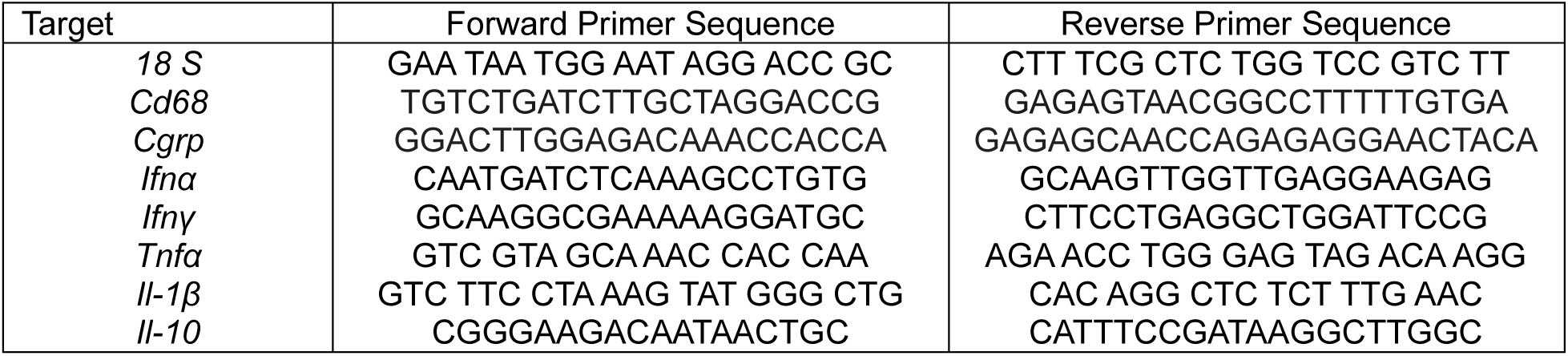
Primer sequences for RT-qPCR.

### Flow Cytometry

Lung tissue was removed from animals without perfusion. Tissues were minced for one minute, then incubated in digestion buffer (5% heat-inactivated fetal bovine serum, 10 mM HEPES, 5 mg/mL liberase, 1:10,000 DNAse I in 1 x HBSS) at 37°C shaking at 225 rpm for 25 minutes until tissue was completely digested. Following digestion, single cell suspensions were prepared by filtration with a 70 μm cell strainer and diluted in PFH buffer (5% heat-inactivated fetal bovine serum, 1 Mm HEPES in 1x Ca2 and Mg2 free DPBS). Cells were treated with hemolysis buffer (0.15M ammonium chloride, 10mM potassium bicarbonate, and 1mM EDTA in diH2O) for 5 minutes at RT, then buffer was quenched using PFH buffer. This suspension was strained using a 40 μm cell strainer. Cells were pelleted by centrifugation, resuspended in PFH buffer, and counted with a hemocytometer. 1 million cells per sample were pelleted by centrifugation then resuspended in 100 μL PFH buffer. Cells were blocked in 1:200 Fc Block (CD16/32) for at least 15 minutes over ice. Cells were then incubated with one of two antibody panels (see tables 3 and 4 below) for 20 minutes over ice covered from light. Cells were pelleted by centrifugation, washed with PFH buffer, pelleted again by centrifugation, then resuspended in 100 μL PFH buffer. Samples were kept covered over ice until analyzed on a FortessaX20 Flow Cytometer (BD Biosciences). The following markers were used to identify lung cell populations: total Myeloid cells (CD45^+^CD11b^+^), Neutrophils (CD45^+^CD11b^+^Ly6G^+^), Monocytes (CD45^+^CD11b^+^Ly6G^-^MHCII^-^Ly6C^+^), Exudate macrophages (CD45^+^CD11b^+^CD64^+^CD24^-^Ly6G^-^Ly6C^+^SiglecF^-^), Alveolar macrophages (CD45^+^CD11b^+^CD64^+^CD24^-^Ly6G^-^Ly6C^-^SiglecF^+^), Dendritic cells (CD45^+^CD11b^+^Ly6G^-^Ly6C^-^CD64^-^CD24^+^CD11c^+^MHCII^+^), Eosinophils (CD45^+^CD11b^+^Ly6G^-^Ly6C^-^CD64^-^CD24^+^CD11c^-^MHCII^-^SiglecF^+^), B cells (CD45^+^TCRβ^-^CD19^+^), T cells (CD45^+^CD19^-^TCRβ^+^), Total CD4 T cells (CD45^+^CD19^-^TCRβ^+^CD4^+^), Naïve CD4 T cells (CD45^+^CD19^-^TCRβ^+^CD4^+^CD62L^+^CD44^-^), Effector CD4 T cells (CD45^+^CD19^-^TCRβ^+^CD4^+^CD62L^-^ CD44^+^), Central Memory CD4 T cells (CD45^+^CD19^-^TCRβ^+^CD4^+^CD62L^+^CD44^+^), Total CD8 T cells (CD45^+^CD19^-^TCRβ^+^CD8^+^), Naïve CD8 T cells (CD45^+^CD19^-^TCRβ^+^CD8^+^CD62L^+^CD44^-^), Effector CD8 T cells (CD45^+^CD19^-^TCRβ^+^CD8^+^CD62L^-^ CD44^+^), Central Memory CD8 T cells (CD45^+^CD19^-^TCRβ^+^CD8^+^CD62L^+^CD44^+^), Natural Killer cells (CD45^+^TCRβ^-^CD19^-^NK1.1^+^). Analyses were performed using fresh, live cells (negative for DAPI staining). Fluorophore compensation was performed using live cells and UltraComp eBeads (Thermo Fisher Scientific). FlowJo software (Treestar) was used for data analyses.

**Table 3.** Flow cytometry antibodies for myeloid <u>cell</u> panel.

| Fluorophore | Antigen | Brand | Catalog Number | Concentration |
| --- | --- | --- | --- | --- |
| BUV395 | CD45 | BD Horizon | 564279 | 1:200 |
| AF700 | I-A/I-E | BioLegend | 107622 | 1:200 |
| BV605 | CD11b | BioLegend | 101257 | 1:200 |
| APC-CY7 | Ly-6G | BioLegend | 127623 | 1:200 |
| APC | CD64 | BioLegend | 139306 | 1:200 |
| PE | CD24 | BioLegend | 101807 | 1:200 |
| PerCP-Cy 5.5 | SiglecF | BD Horizon | 565526 | 1:200 |
| BV785 | CD11c | BioLegend | 117336 | 1:200 |
| PE-CY7 | LY6C | BioLegend | 128017 | 1:200 |

**Table 4.** Flow cytometry antibodies for lymphocyte panel.

| Fluorophore | Antigen | Brand | Catalog Number | Concentration |
| --- | --- | --- | --- | --- |
| BUV395 | CD45 | BD Horizon | 564279 | 1:200 |
| PE-CY7 | NK1.1 | BioLegend | 108714 | 1:200 |
| PE | TCRB | BioLegend | 109207 | 1:200 |
| BV605 | CD4 | BioLegend | 100451 | 1:200 |
| BV785 | CD8 | BioLegend | 100750 | 1:200 |
| APC | CD19 | BioLegend | 115511 | 1:200 |
| PE-Cy5 | CD62L | BioLegend | 104410 | 1:200 |

### Statistics

Statistical tests were performed using GraphPad Prism 9. Asterisks denote the following significance thresholds: n.s. for p > 0.05, * for p ≤ 0.05, ** for p ≤ 0.01, *** for p ≤ 0.001. Unless otherwise noted, data plots display mean values ± S.E.M.

## ACKNOWLEDGEMENTS

This work was supported by funding from NIH grant F31NS134281.

Influenza A viral stocks were provided by the lab of Dr. Nicholas Heaton. Special thanks to Brook Heaton, Cait Hamele, Andy Miranda, and Ariel Sprurrier for their guidance and assistance with pulmonary procedures including viral titer measurement, viral infection, viral handling, and pulmonary tissue handling.

Figurea 1, 3, and S3 include graphics made using BioRender. Confirmation of the author’s publication and licensing rights and are available to view online for Figure 1A (https://BioRender.com/6q25un1), 3A (https://BioRender.com/78lhiwq), and S3A (https://BioRender.com/yqmirl0).

Special thanks to Francesco Paolo Ulloa Severino for assistance with analysis of influenza A infection response mapping in the brain during preliminary experiments.

Special thanks to Mari Shinohara, Estefany Reyes, and Miranda Lumbreas for their guidance and assistance with flow cytometry experiments.

The authors have no conflicting interests.

## FIGURE CAPTIONS

**Figure S1.**
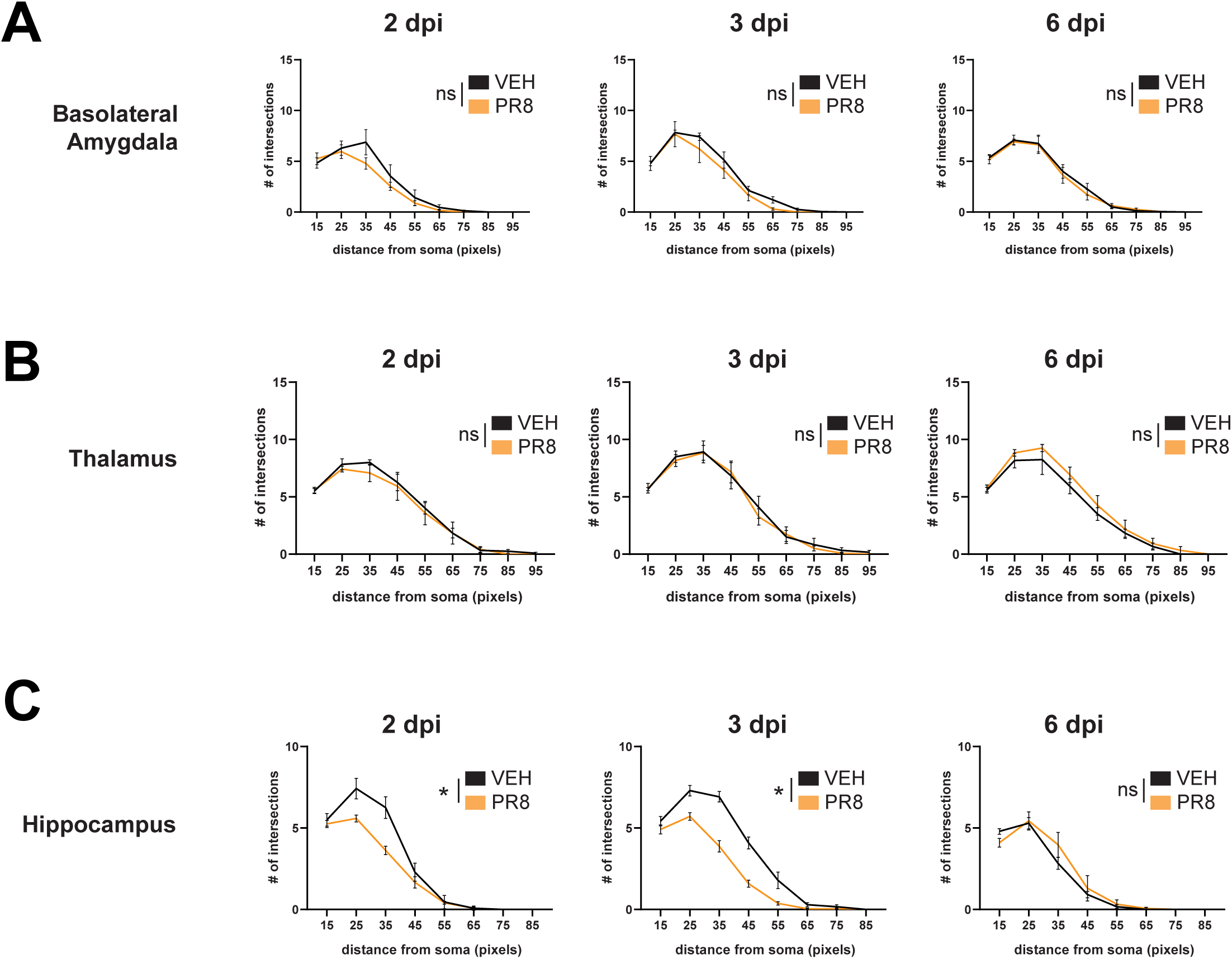
(A) Quantification of microglia morphology in the basolateral amygdala at 2, 3, and 6 dpi by Sholl analysis [2-way ANOVA, n = 4 – 5]. (B) Quantification of microglia morphology in the thalamus at 2, 3, and 6 dpi by Sholl analysis [2-way ANOVA, n = 4 – 5]. (C) Quantification of microglia morphology in the hippocampus at 2, 3, and 6 dpi by Sholl analysis [2-way ANOVA, n = 4 – 5].

**Figure S2.**
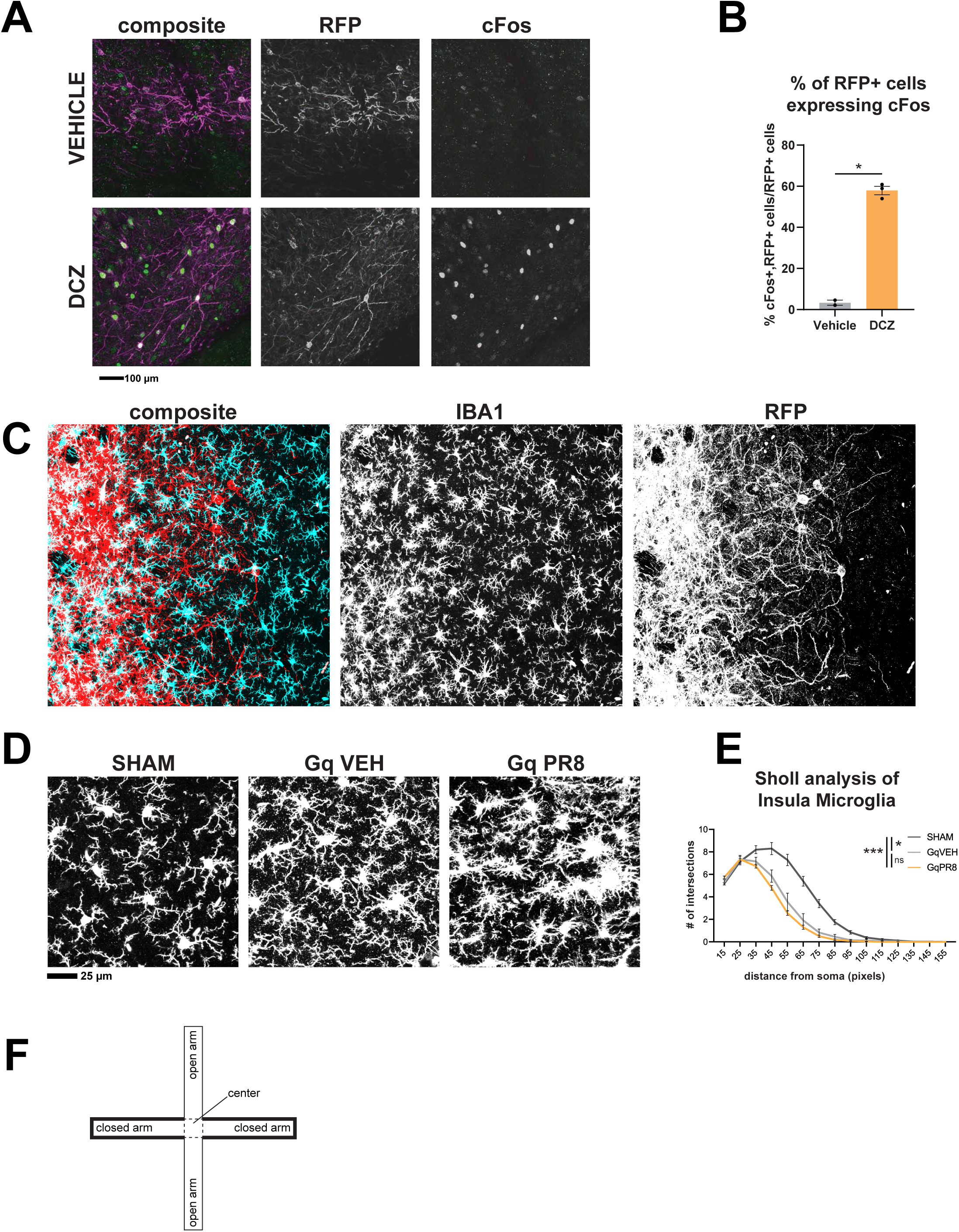
(A) Representative immunofluorescence images of RFP and cFos staining in brain tissue of animals receiving stereotaxic injection of cre-dependent hM3Dq plasmid collected at 90 minutes after i.p. injection of DCZ or vehicle solution. (B) Quantification of RFP+, cFos+ (double-positive) cells as a fraction of all RFP+ cells [T-test, n = 2 – 3]. (C) Representative immunofluorescence image from the posterior insula of a Gq-VEH animal demonstrating microglia morphology (IBA1) changes after DCZ stimulation in relation to captured neuron (RFP) density. (D-E) Representative immunofluorescence images of IBA1 signal in the posterior insula collected 115 minutes after DCZ stimulation (E) Quantification of microglia morphology in the posterior insula [1-way ANOVA, n = 6 – 9]. (F) Schematic of elevated plus maze arena showing closed arms, open arms, and center.

**Figure S3.**
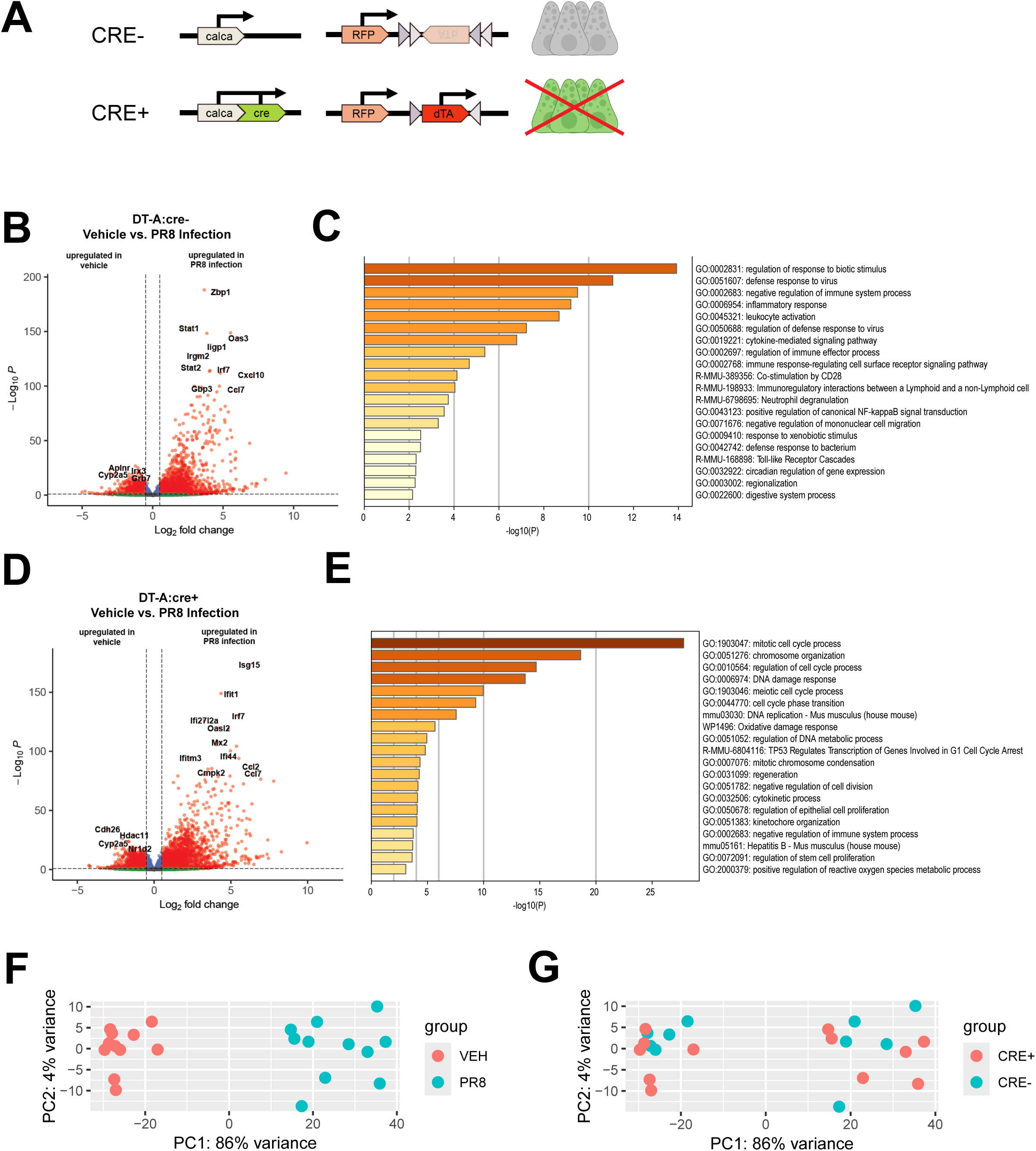
(A) Schematic representation of floxed gene expression and PNEC ablation in cre+ but not in cre- animals. (B) Volcano plot showing DEGs in lung homogenate of DT-A:cre- animals given PR8 infection or vehicle treatment at 6 dpi; significant DEGs are shown in red [n = 5 – 6]. (C) Gene ontology of DEGs in lung homogenate of DT-A:cre- animals given PR8 infection or vehicle treatment at 6 dpi [n = 5 – 6]. (D) Volcano plot showing DEGs in lung homogenate of DT-A:cre+ animals given PR8 infection or vehicle treatment at 6 dpi; significant DEGs are shown in red [n = 5 – 6]. (E) Gene ontology of DEGs in lung homogenate of DT-A:cre+ animals given PR8 infection or vehicle treatment at 6 dpi [n = 5 – 6]. (F-G) PCA plot of bulk sequencing data from whole lung homogenate of DT-A:cre+ and DT-A:cre- animals given PR8 infection or vehicle treatment colored by infection status (F) or genotype (G).

**Figure S4.**
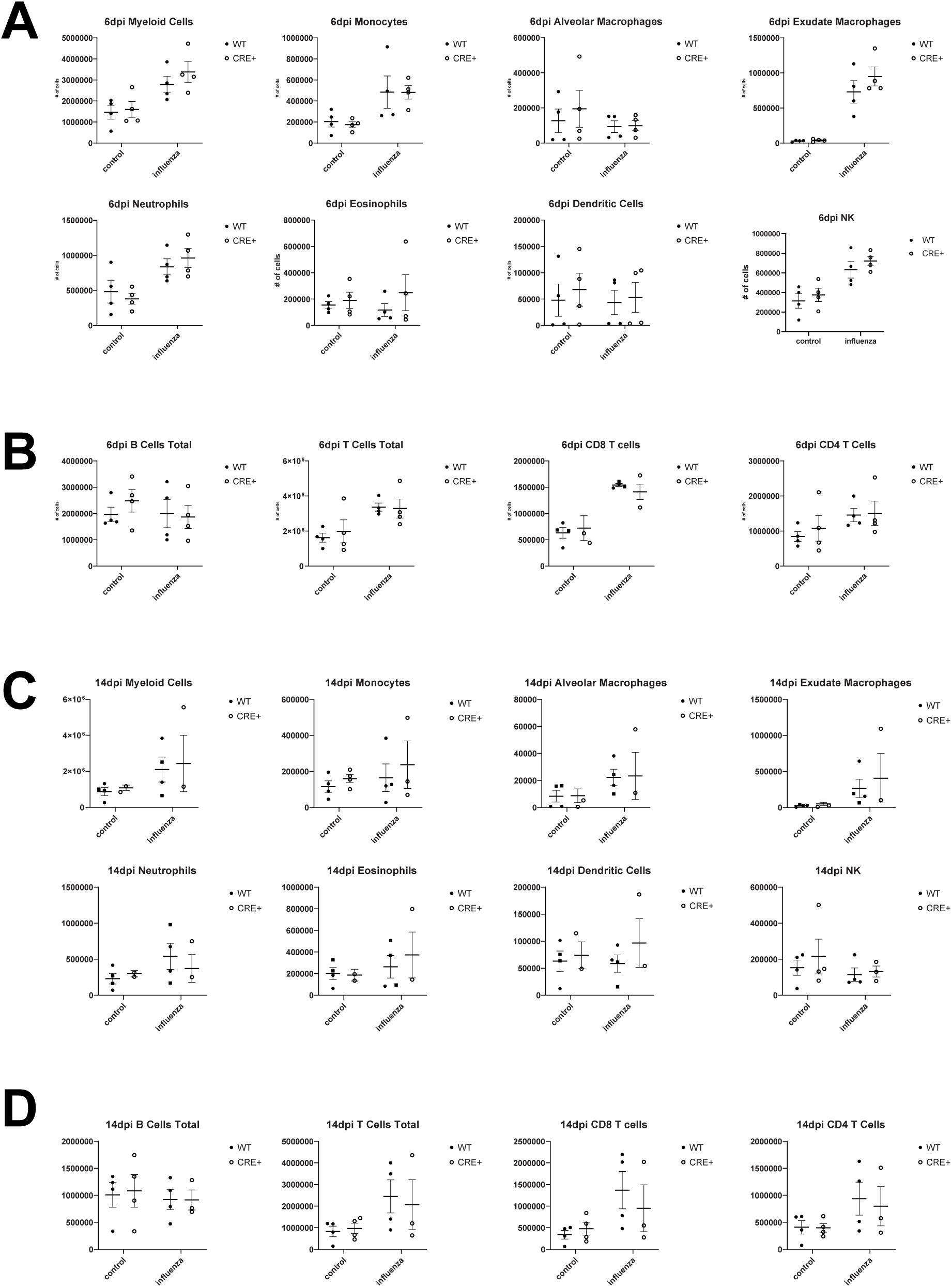
(A-B) Box plots counting myeloid cell populations (A) and lymphocyte cell populations (B) measured by flow cytometry at 6 dpi in dissociated whole lung [2-way ANOVA, n = 4]. (C-D) Box plots counting myeloid cell populations (C) and lymphocyte cell populations (D) measured by flow cytometry at 14 dpi in dissociated whole lung [2-way ANOVA, n = 4].

**Figure S5.**
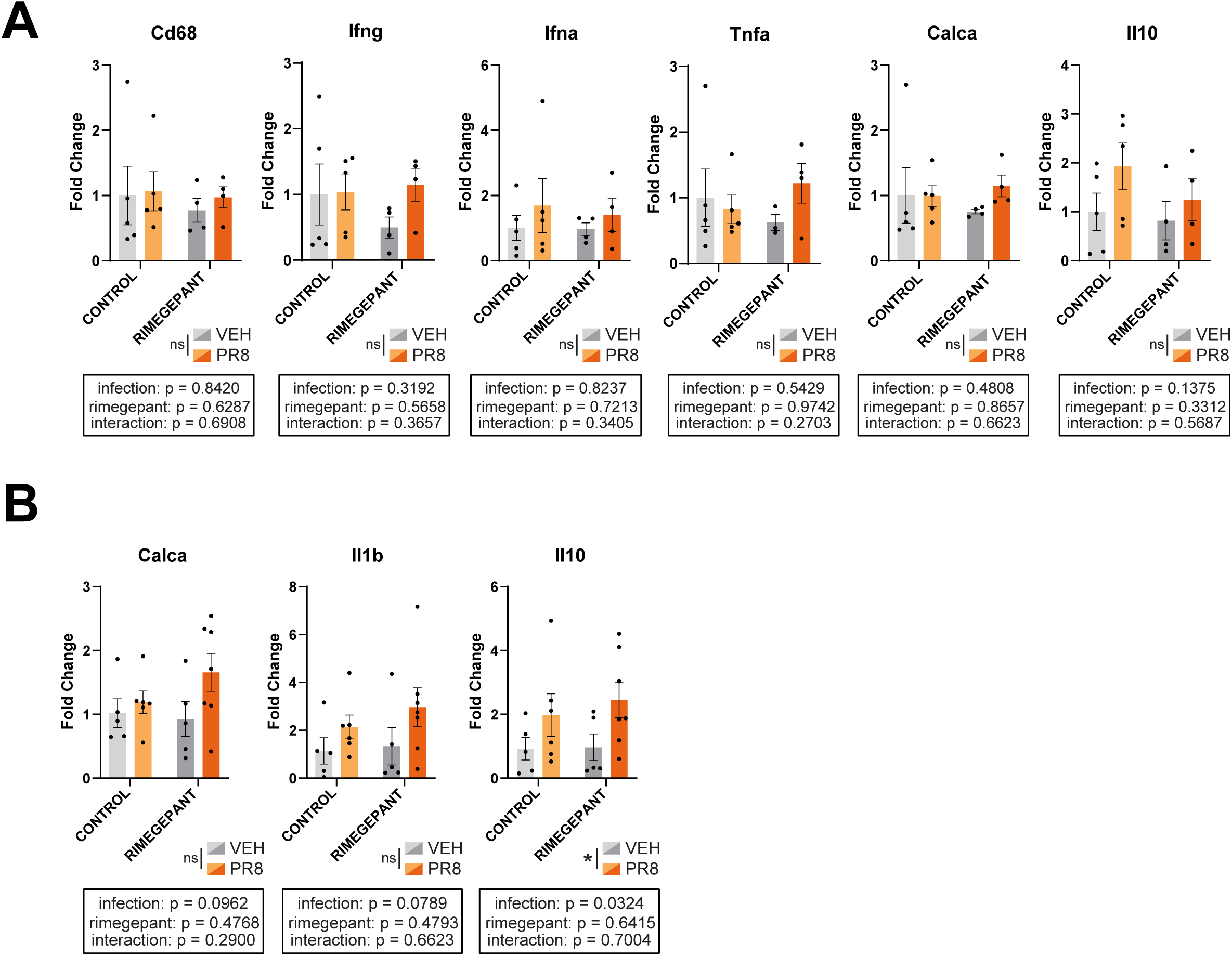
(A) Fold change (ΔΔC_T_) of RNA expression for genes measured by RT-qPCR [2-way ANOVA, n = 4 – 5] in lung homogenate after PR8 infection or vehicle treatment and rimegepant or vehicle control injection. (B) Fold change (ΔΔC_T_) of RNA expression for genes measured by RT-qPCR [2-way ANOVA, n = 5 – 7] in lung homogenate after 3 days of DCZ stimulation.

